# A Genome Model Linking Birth Defects to Infections

**DOI:** 10.1101/674093

**Authors:** Bernard Friedenson

## Abstract

The purpose of this study was to test the hypothesis that infections are linked to chromosomal anomalies that cause neurodevelopmental disorders. In children with disorders in the development of their nervous systems, chromosome anomalies known to cause these disorders were compared to microbial DNA, including known teratogens. Genes essential for neurons, lymphatic drainage, immunity, circulation, angiogenesis, cell barriers, structure, epigenetic and chromatin modifications were all found close together in polyfunctional clusters that were deleted or rearranged in neurodevelopmental disorders. In some patients, epigenetic driver mutations also changed access to large chromosome segments. These changes account for immune, circulatory, and structural deficits that accompany neurologic deficits. Specific and repetitive human DNA encompassing large deletions matched infections and passed rigorous artifact tests. Deletions of up to millions of bases accompanied infection-matching sequences and caused massive changes in the homologies to foreign DNAs. In data from three independent studies of private, familial and recurrent chromosomal rearrangements, massive changes in homologous microbiomes were found and may drive rearrangements and encourage pathogens. At least one chromosomal anomaly was found to consist of human DNA fragments with a gap that corresponded to a piece of integrated foreign DNA. Microbial DNAs that match repetitive or specific human DNA segments are thus proposed to interfere with the epigenome and highly active recombination during meiosis, driven by massive changes in the homologous microbiome. Abnormal recombination in gametes produces zygotes containing rare chromosome anomalies which cause neurologic disorders and non-neurologic signs. Neurodevelopmental disorders may be examples of assault on the human genome by foreign DNA at a critical stage. Some infections may be more likely tolerated because they resemble human DNA segments. Further tests of this model await new technology.

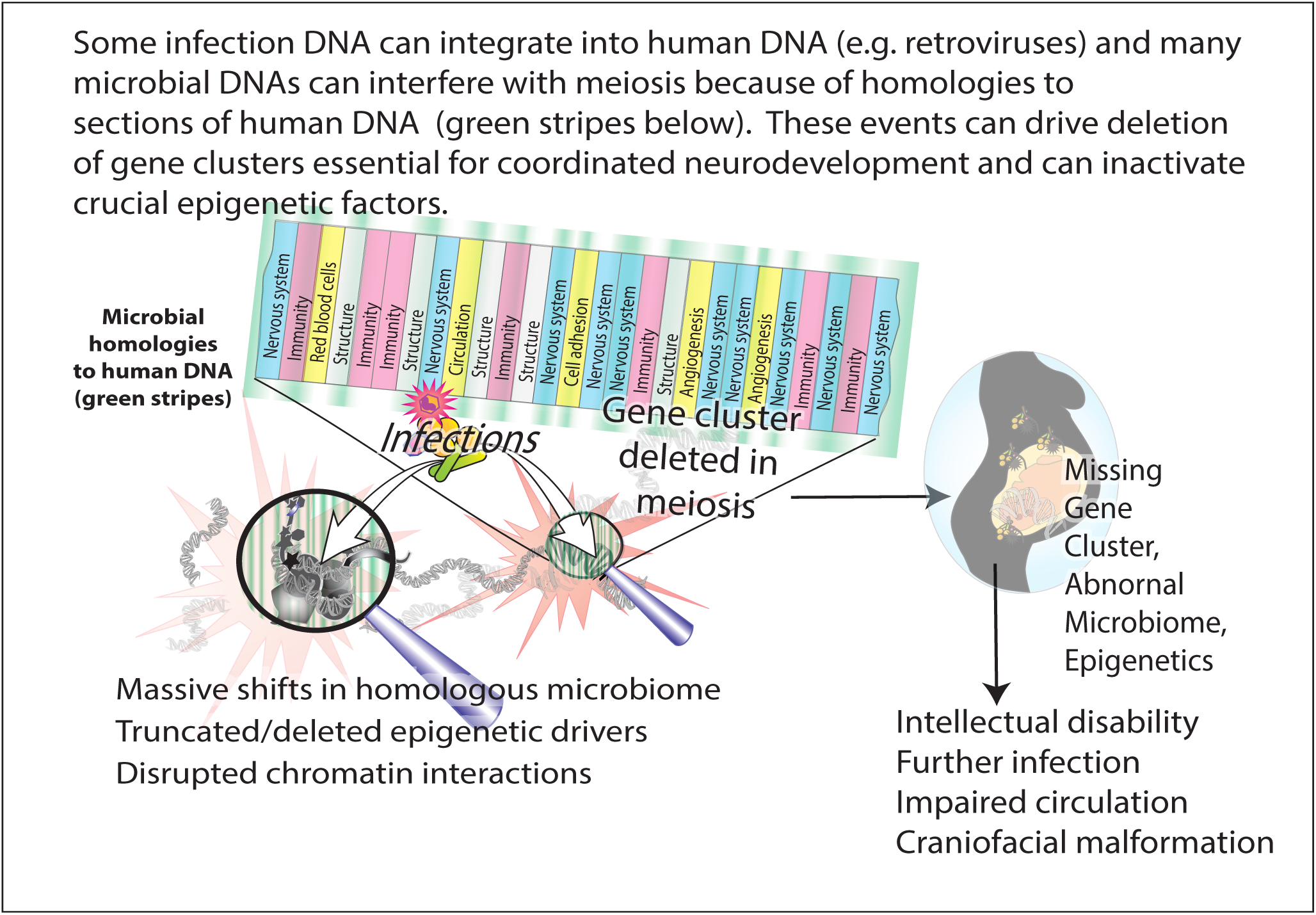

## Introduction

An approach to preventing neurodevelopmental disorders is to gain better understanding of how neurodevelopment is coordinated and then to identify interference from environmental, genomic and epigenomic factors.The development of the nervous system requires tight regulation and coordination of multiple functions essential to protect and nourish neurons. As the nervous system develops, the immune system, the circulatory system, cranial and skeletal systems must all undergo synchronized and coordinated development. Neurodevelopmental disorders follow the disruption of this coordination.

A significant advance in genome sequence level resolution of balanced cytogenetic abnormalities greatly improves the ability to document changes in regulation and dosage for genes critical to the function of the neurologic system. Based on DNA sequence analyses, some chromosome rearrangements have been identified as causing individual congenital disorders because they disrupt genes essential for normal development ^1–3^. There is poor understanding and no effective treatment for many of these overwhelming abnormalities. Signs and symptoms include autism, microcephaly, macrocephaly, behavioral problems, intellectual disability, tantrums, seizures, respiratory problems, spasticity, heart problems, hearing loss, and hallucinations ^1^. Because the abnormalities do not correlate well with the outcome, genetic counseling is difficult and uncertain ^3^.

In congenital neurologic disease, inheritance is usually autosomal dominant and the same chromosomal abnormalities occur in every cell.The genetic events that lead to most neurodevelopmental disorders are not understood ^4^ but several maternal infections and other lifestyle factors are known to interfere.

DNA homology between microbes and humans is a known fact and DNA swapping between vertebrates and invertebrates has been reported. An early draft of the human genome found human genomes have undergone lateral gene transfer to incorporate microorganism genes ^5^. Lateral transfer from bacteria may have generated many candidate human genes ^6^. Genome-wide analyses in animals found up to hundreds of active genes generated by horizontal gene transfer. Fruit flies and nematodes have continually acquired foreign genes as they evolved. Although these transfers are thought to be rarer in primates and humans, at least 33 previously unreported examples of horizontally acquired genes were found ^6^. These findings argue that horizontal gene transfer continues to occur to a larger extent than previously thought. Transferred genes that survive have been largely concerned with metabolism and make important contributions to increasing biochemical diversity^7^.

The present work implicates foreign DNA largely from infections as a cause of the chromosome anomalies that cause birth defects. Infections replicate within the human CNS by taking advantage of immune deficiencies such as those traced back to deficient microRNA production ^8^ or other gene losses. Disseminated infections can then interfere with the highly active DNA break repair process required during meiosis. The generation of gametes by meiosis is the most active period of recombination which occurs at positions enriched in epigenetic marks on chromatin. Hundreds of double strand breaks accompany meiotic recombination^9^. Gametes with errors in how this recombination occrs cause chromosome anomalies in the zygote. In contrast to oocytes, meiotic recombination in sperm cells occurs continuously after puberty.

The exact DNA sequences of known pathogenic rearrangements in individual, familial and recurrent congenital disorders ^1–3^ make it possible to test for association with foreign DNA. Even rare developmental disorders can be screened for homology to infections within altered epigenomes and chromatin structures. This screening may assist counseling, diagnosis, prevention, and early intervention.

The results showed that DNA abnormalities in some neurodevelopmental disorders closely match DNA in multiple infections that extend over long linear stretches of human DNA and often resemble repetitive human DNA sequences. Massive changes in the identity and distribution of sets of homologous microorganisms that accompany chromosome abnormalities may drive and stabilize them. Removing competition by changing the normal microbiome may encourage pathogens.

The affected sequences are shown to exist as linear clusters of genes closely spaced in two dimensions. Interference from infection can also delete or damage human gene clusters and change epigenetic functions that coordinate neurodevelopment. This microbial interference accounts for immune, circulatory, and structural deficits that accompany neurologic deficits.

Congenital neurodevelopmental disorders are thus viewed as resulting from an assault on human DNA by foreign DNA and perhaps selection of infecting microorganisms based on their similarity to host DNA. It is important to remember that effects in two dimensions can sometimes alter 3D topology as well ^10^.

Testing and verifying predictions from a viable model may spur the development of methods for identifying contributions from infections in intractable rare disorders that are not now available. Convergent arguments from testing predictions based on any proposed model might lessen effects of limitations in currently available technology.

## Materials and Methods

### Data Sources

DNA sequences from acquired congenital disorders were from published whole genome sequences at chromosome breakpoints and rearrangement sites ^1–3^. Comparison to multiple databases of microbial sequences determined whether there was significant human homology. Emphasis was on cases with strong evidence that a particular human chromosome rearrangement was pathologic for the congenital disorder. Patients in the 3 major studies ^1–3^ used in this analysis were 98 females and 144 males. Of the ages most, most (46) patients were under age 10, 7 were age 10-20, 6 were age 20-40 and 1 patient was older than 40.

### Testing for homology to microbial sequences

Hundreds of different private rearrangements in patients with different acquired congenital disorders were tested for homology ^11^ against non-human sequences from microorganisms known to infect humans as follows: Viruses (taxid: 10239), and retroviruses including HIV-1 (taxid:11676), human endogenous retroviruses (Taxids:45617, 87786, 11745, 135201, 166122, 228277and 35268); bacteria (taxid:2); Mycobacteria (taxid:85007); fungi (taxid; 4751), and chlamydias (taxid:51291)

Because homologies represent interspecies similarities,“Discontinuous Megablast” was most frequently used, but long sequences were sometimes tested against highly similar microbial sequences. Significant homology (indicated by homology score) occurs when microbial and human DNA sequences have more similarity than expected by chance (E value<=e-10) ^12^. Confirmation of microorganism homologies was done by testing multiple variants of complete microorganism genomes against human genomes and by extending analyses to 20000 matches.

Various literature analyses have placed Alu repeats into 8 subfamilies having consensus sequences (GenBank; accession numbers U14567 - U14574). Microbial sequences were independently compared to all 8 consensus Alu sequences and to 442 individual AluY sequences.

### Chromosome localizations

The positions of microbial homologies in human chromosomes were determined using BLAT or BLAST. Comparisons were also made to cDNAs based on 107,186 Reference Sequence (RefSeq) RNAs derived from the genome sequence with varying levels of transcript or protein homology support. Tests for contamination by vector sequences in these non-templated sequences were also carried out with the BLAST program. Inserted sequences were also compared to Mus musculus GRCM38.p4 [GCF_000001635.24] chromosome plus unplaced and unlocalized scaffolds (reference assembly in Annotation Release 106). Homology of inserted sequences to each other was tested using the Needleman and Wunsch algorithm. Lists of total homology scores for microbes vs human chromosome rearrangements were compared by the Mann-Whitney U test using the StatsDirect Statistical analysis program. Fishers test was also used.

## Results

### Interdependent functions are clustered together on the same chromosome segment

The nervous system has a close relationship to structures essential for immunity, circulation, cell barriers and protective enclosures. In chromosome segments deleted in neurologic disorders, genes essential for all these functions must develop in concert. Genes for these related functions are located close to each other on the same linear segment of a chromosome (Fig. 1). Deletions at 4q34 in patient DGAP161 are shown in Fig. 1. The genes within the 4q34 deletion in each of four categories tested are color coded in Fig.1. Fig 1 is representative of six other chromosome bands that were also tested and gave similar results: 2q24.3; 6q13-6q14.1; 10p14-10p15.1; 13q14.2; 18p11.22-p11.32; and 19q12-q13.1. Deletion of these clustered arrangements has been correlated with serious neurodevelopmental disorders^1^. Alternatively-spliced forms of the same gene may encode for pleiotropic functions that must be synchronized and coordinated among diverse cells. Multiple functions for the same gene in different cell types are commonly found. Hormonal signaling represents a major control mechanism^13^.

**Fig. 1.**
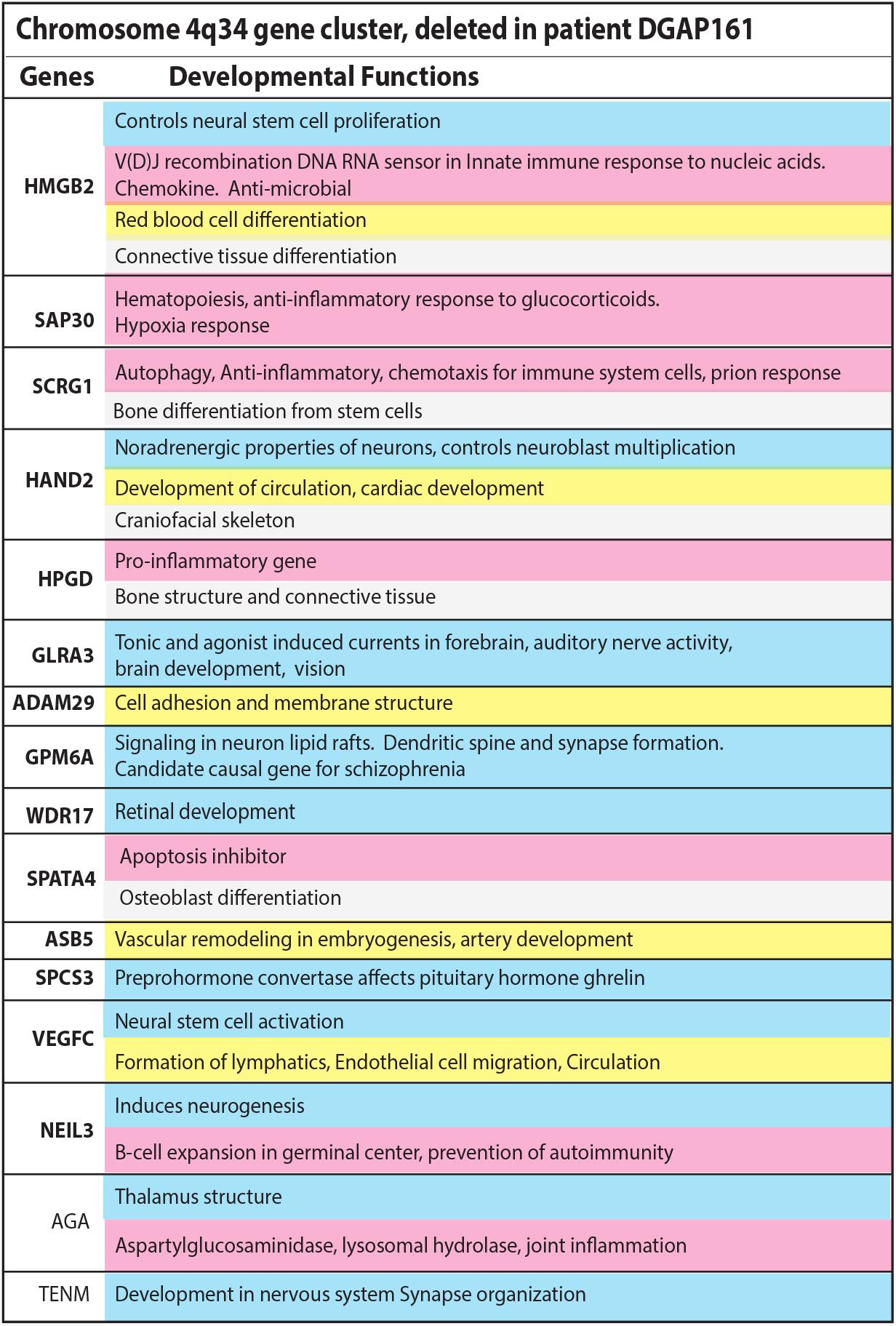
Chromosome 4q34 as a typical example of close relationships between nervous system genes and genes for other essential developmental functions. Pleiotropic genes for multiple interdependent systems appear in clusters on the same chromosomes. Nervous system genes are near genes essential for the immune system, connections to lymphatic circulation, ability to form tight junctions, structural enclosures and chromatin control. Clusters of genes encoding these and other interdependent functions on chromosome segments are deleted in private neurodevelopmental disorders. These losses increase susceptibility to infections that have DNA homologous to long stretches of human DNA. In the example shown, genes are listed in the order they occur on 4q34. Developmental functions are to the right of the gene symbol. Blue genes are associated with the nervous system, pink with the immune system. Yellow genes have functions associated with angiogenesis, or lymphangiogenesis or cell barriers. Genes for development of essential bone structure or connective tissues needed to protect and house the nervous system are light grey. Isoforms of the same gene may encode different functions in different cellular locations. Consistent results were obtained from deletions involving 6 other chromosome bands (see text).

### Many deleted gene clusters include long stretches of DNA strongly related to infections

To investigate the chromosomal segment deletions that likely cause neurodevelopmental disorders, homologies to infection were tested in sequences within and flanking deleted clusters. Strong homologies to infections were interspersed. To demonstrate the extent of these relationships, deleted 4q34 chromosome segment (Patient DGAP161 ^1^) was tested for homology to microbes in 200 kb chunks.

Fig. 2 shows that stretches of homology to microorganisms are distributed throughout chromosome 4q34. Only 10 homologous microbes are shown for each 200 kb division, but there are up to hundreds, giving a total of many thousands of potentially homologous microorganisms throughout the 4q34 deletion.

**Fig. 2.**
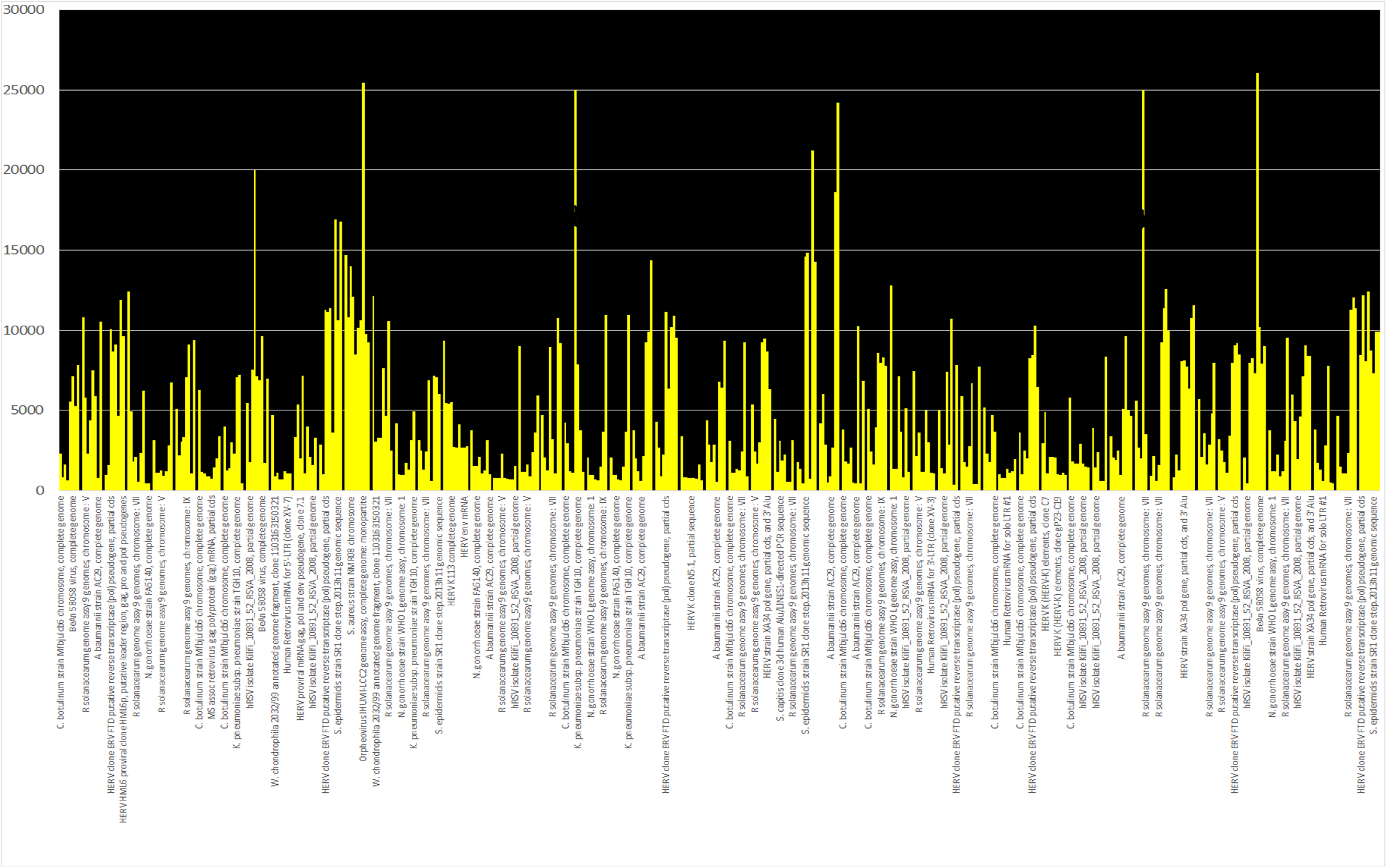
Example of total homologies to microbial sequences dispersed throughout the normal 4q34 chromosome segment deleted in patient DGAP161. The top 10 homologies are shown for each 200,000 bp segment and are listed within each of these 57 segments in order of their scores. The Y axis plots the total homology scores and the x axis lists the individual microorganism. The mean E value was 5.3e-90 (range 0.0 to 1e-87). F. tularensis had values of 33,368 and 79,059 which were truncated at 25000 to show more detail for the other microbial matches.

### Enormous effects on microbial homologies accompany changes in junction sequences between chromosome bands caused by deletions

Fig 3 represents a snapshot of how local microbial homologies shift when a chromosome segment is deleted. Fig. 3 (top) represents the normal 4q33-4q34 junction and its change after the 4q34 deletion to the new junction 4q33-4q35. Around the breakpoint and especially 3’ to it, microbial homologues become very different. Although the representations of microbial homologies as red rectangles appear to be small on Fig. 3 (top), the matching sequences actually extend for hundreds of base pairs. Normally, there are multiple homologies to a variety of microorganisms at the breakpoint (yellow rectangle) which may help destabilize the area. After the deletion, the junction formed now has new and strong homology to pathogens N. gonorrhoeae, k. pneumoniae, clostridium botulinum and other potential pathogens. Homology to some isolates of γ-herpesvirus 4 (EBV) extends into the newly juxtaposed 4q35 region. The deletion changes the sets of homologous microorganisms (Mann Whitney test P=0.0089). Both maximum or total homology scores show that microbial distributions become very different after the rearrangement (P=0.0178). Extendng these analyses to test for 20,000 homologues further supports this conclusion (data not shown). Depending on which microbes exist in the microenvironment, the changes in the microbiome could easily cause a change in free energies to drive, direct and guide the chromosome rearrangement. Changes in topological chromosome interactions are also likely.

**Fig. 3.**
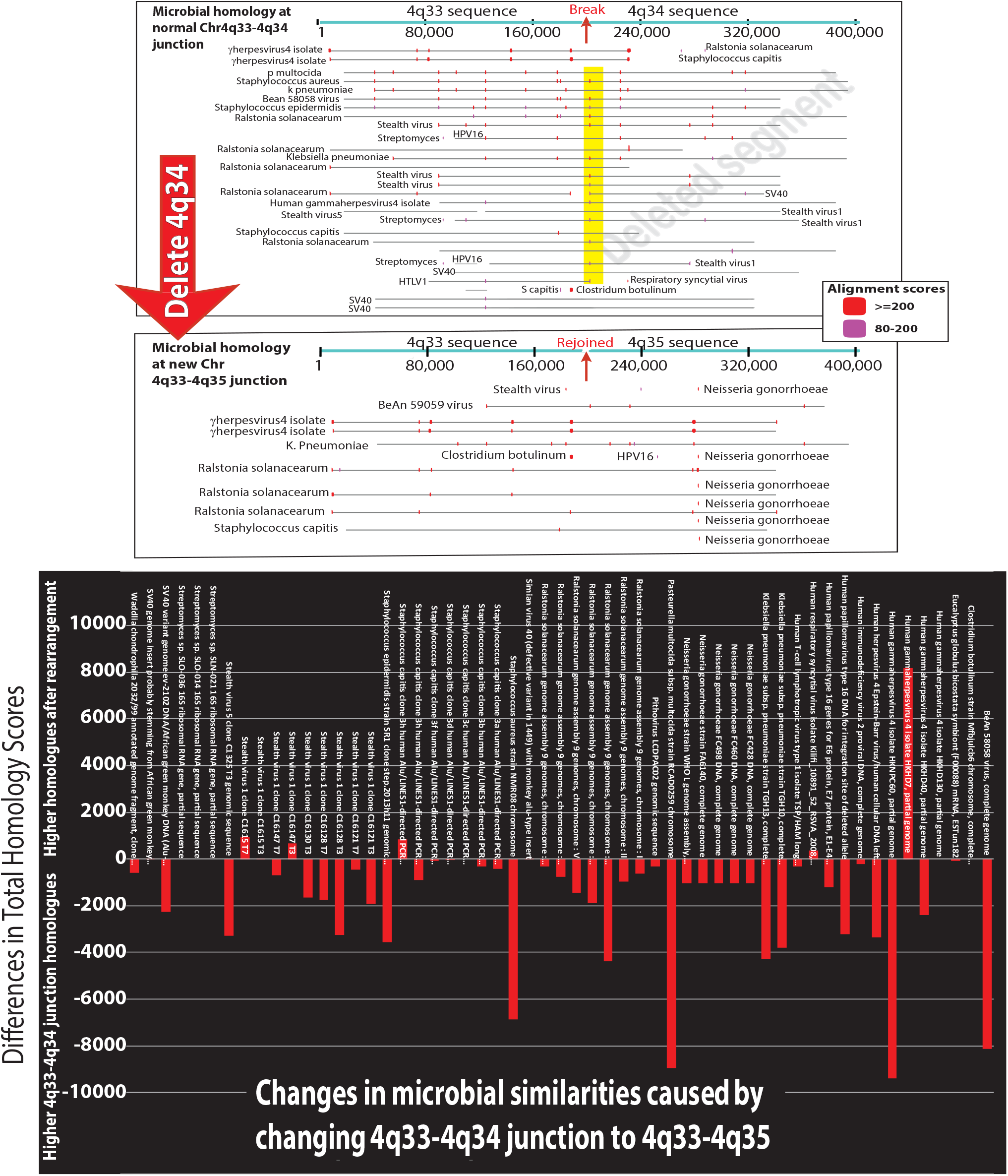
Snapshot of very large differences in local microbial homologies in one 200 kb section of chromosome 4q34. The same 200 kbp sections after vs before deletion of 4q34 are shown. Top: The 4q34 deletion changes the local microbiome homologous to human sequences in this region. The top panel shows the breakpoint of the deletion. There is homology to stealth viruses and bacteria near the breakpoint (yellow box). After the deletion occurs, homology to human gammaherpes virus4 (EBV) extends into 4q35 and there is now homology to pathogens: clostridium, k. pneumoniae and N. gonorrhoeae. Bottom: In the rearranged 200 kb segment, very different microorganism DNAs have homology to the human chromosome segment and are shown above the x-axis. Increasing the testing to include 20,000 sequences gave the same conclusion.

### Deleted segments in familial chromosome anomalies point toward a general mechanism for infection as a cause of neurodevelopmental disorders

Genome sequencing of the entire family may be necessary ^3^ because some family members carry balanced chromosomal translocations but do not have neurodevelopmental disease. In three of four families with familial balanced chromosomal translocations, patient specific unbalanced deletions were found but the results did not overlap any database of human reference genomes ^3^. A disease associated deletion in the study of Aristidou et al. (Family 2)^3^ was tested by comparing equivalent numbers of bps at the junction sequence created by the deletion vs the original chromosome junction sequence without the deletion (GRCh37:Chr16:49,741,265-49,760,865).

In Fig 4, the changes are enormous, involving new distributions (Mann-Whitney P<0.0001) and different homologous microorganisms. Microbial homologies were confined to much more limited areas of human DNA and their overall complexity in the rearranged human sequences decreases. Some microorganisms newly found in the rearranged sequence are known human pathogens. Their presence emphasizes how different the local microbiome has become. This result was run many times and results were consistent if corrected or uncorrected (result shown) for low complexity human sequences and even if the test sets of sequences were varied somewhat. In agreement with results presented later in Table 1, critical genes linked to epigenetic modifications were among those interrupted by cryptic rearrangements.

**Table 1.**
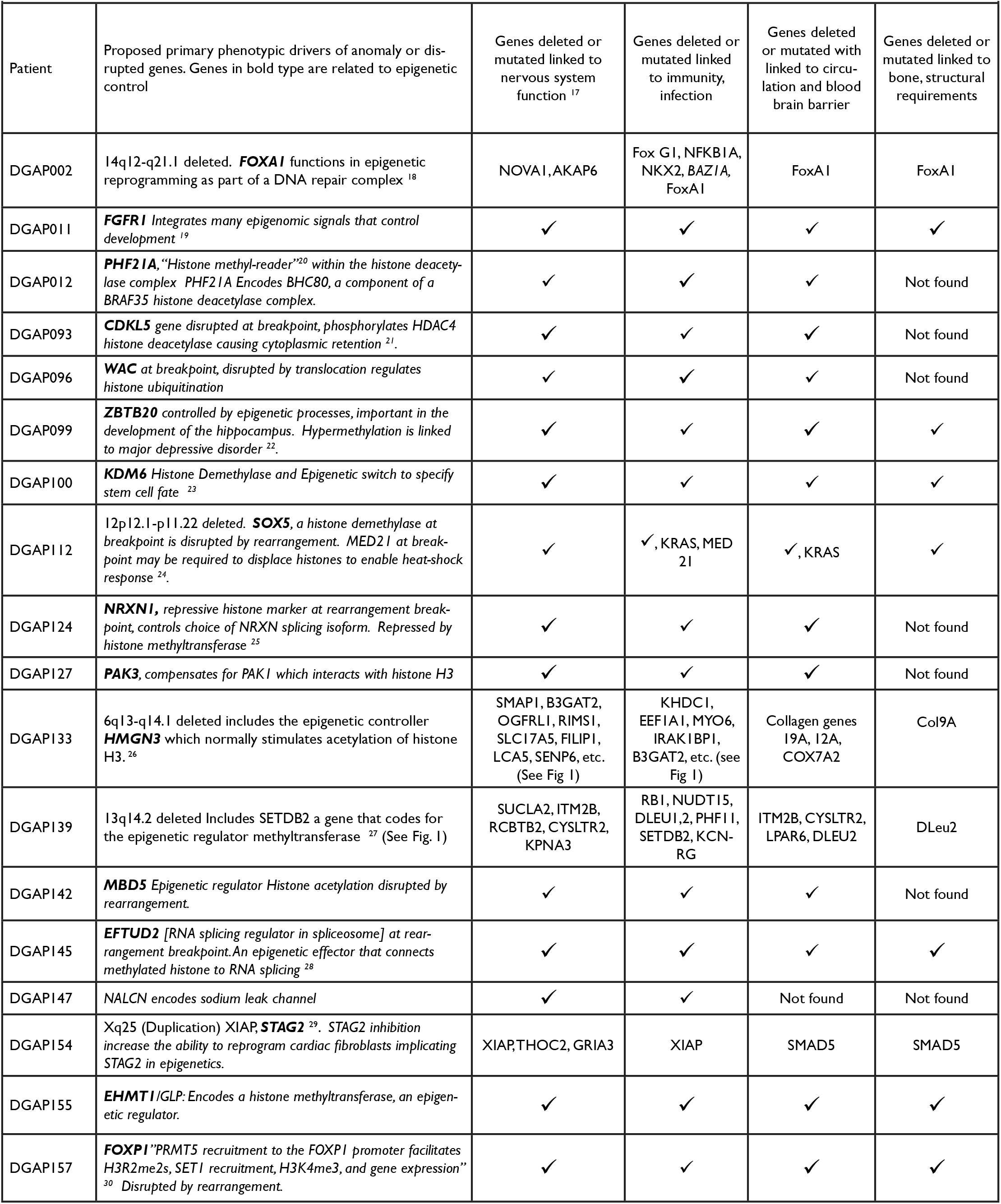
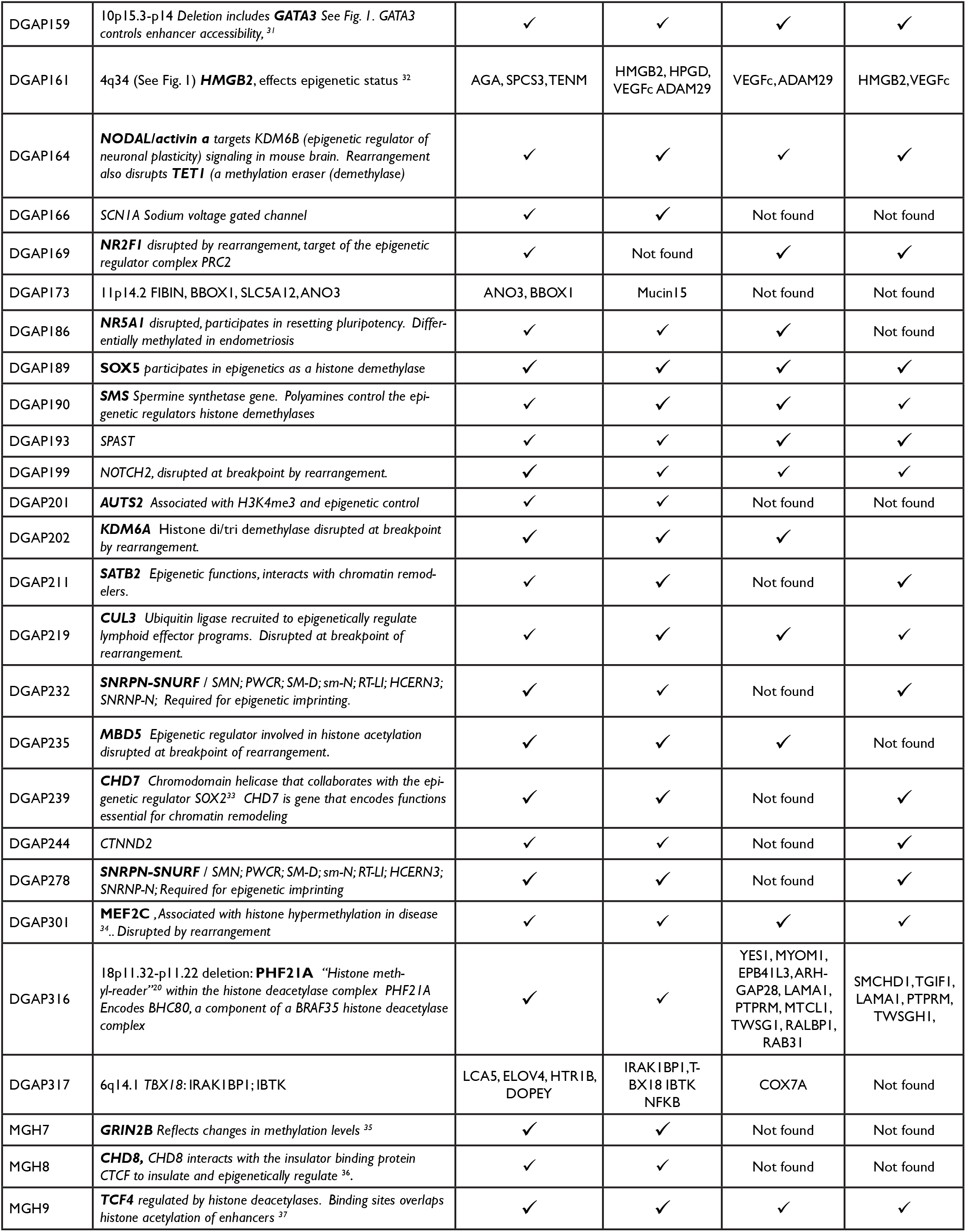
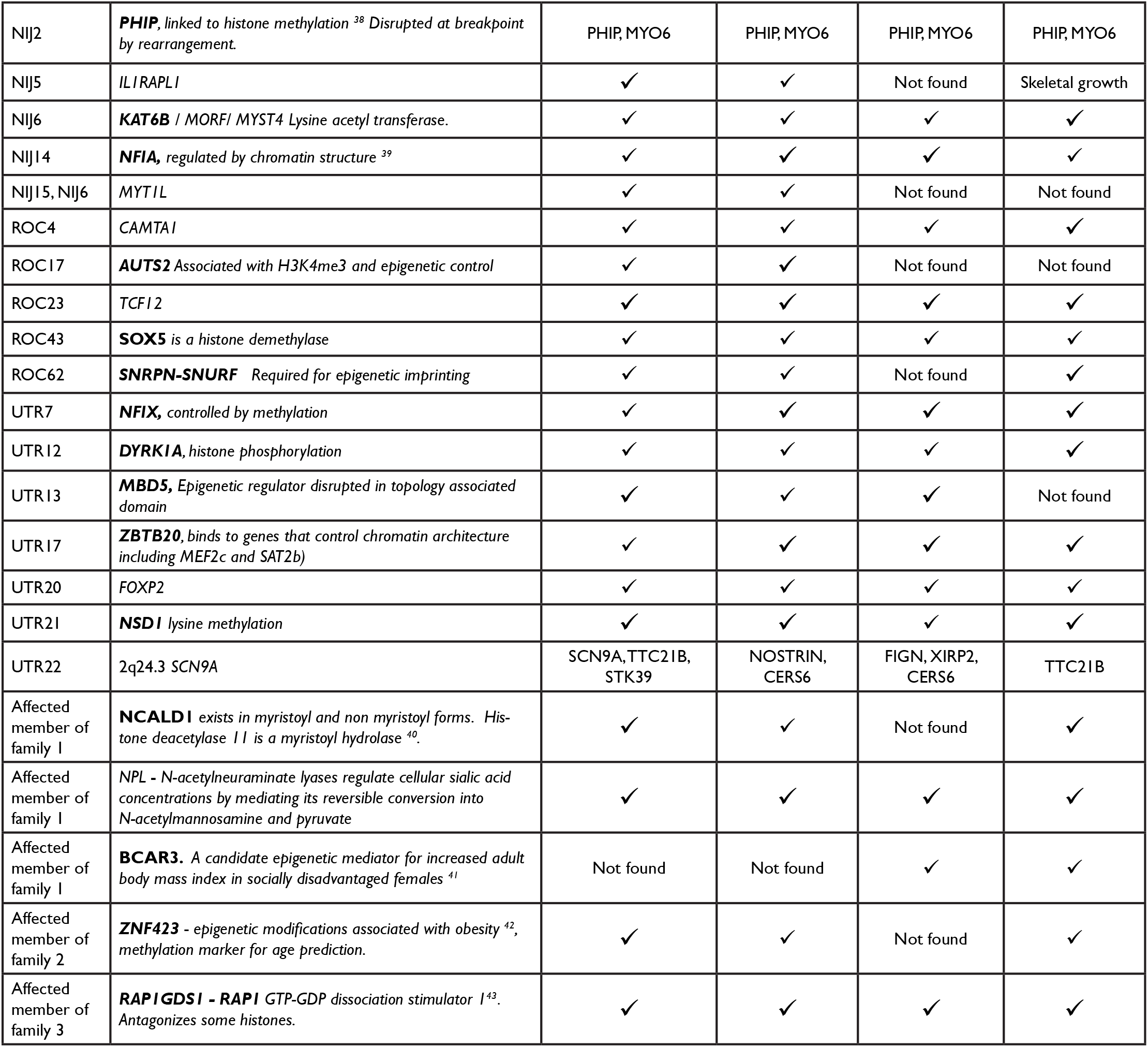
Epigenetic functions of mutated and deleted genes in neurodevelopmental disorders relate neurologic deficits to deficits in the immune system, the circulatory system and structural genes. Checkmark indicates identical gene with mainly epigenetic functions is listed in column 2. Other genes that do not completely match column 2 are listed individually.

| Patient | Proposed primary phenotypic drivers of anomaly or disrupted genes. Genes in bold type are related to epigenetic control | Genes deleted or mutated linked to nervous system function <sup>17</sup> | Genes deleted or mutated linked to immunity, infection | Genes deleted or mutated with linked to circulation and blood brain barrier | Genes deleted or mutated linked to bone, structural requirements |
| --- | --- | --- | --- | --- | --- |
| DGAP002 | 14q12-q21.1 deleted. **FOXA1** functions in epigenetic reprogramming as part of a DNA repair complex <sup>18</sup> | NOVA1, AKAP6 | Fox G1, NFKB1A, NKX2, BAZ1A, FoxA1 | FoxA1 | FoxA1 |
| DGAP011 | **FGFR1** Integrates many epigenomic signals that control development <sup>19</sup> | ✓ | ✓ | ✓ | ✓ |
| DGAP012 | **PHF21A**, "Histone methyl-reader"<sup>20</sup> within the histone deacetylase complex PHF21A Encodes BHC80, a component of a BRAF35 histone deacetylase complex. | ✓ | ✓ | ✓ | Not found |
| DGAP093 | **CDKL5** gene disrupted at breakpoint, phosphorylates HDAC4 histone deacetylase causing cytoplasmic retention <sup>21</sup>. | ✓ | ✓ | ✓ | Not found |
| DGAP096 | **WAC** at breakpoint, disrupted by translocation regulates histone ubiquitination | ✓ | ✓ | ✓ | Not found |
| DGAP099 | **ZBTB20** controlled by epigenetic processes, important in the development of the hippocampus. Hypermethylation is linked to major depressive disorder <sup>22</sup>. | ✓ | ✓ | ✓ | ✓ |
| DGAP100 | **KDM6** Histone Demethylase and Epigenetic switch to specify stem cell fate <sup>23</sup> | ✓ | ✓ | ✓ | ✓ |
| DGAP112 | 12p12.1-p11.22 deleted. **SOX5**, a histone demethylase at breakpoint is disrupted by rearrangement. MED21 at breakpoint may be required to displace histones to enable heat-shock response <sup>24</sup>. | ✓ | ✓, KRAS, MED21 | ✓, KRAS | ✓ |
| DGAP124 | **NRXN1**, repressive histone marker at rearrangement breakpoint, controls choice of NRXN splicing isoform. Repressed by histone methyltransferase <sup>25</sup> | ✓ | ✓ | ✓ | Not found |
| DGAP127 | **PAK3**, compensates for PAK1 which interacts with histone H3 | ✓ | ✓ | ✓ | Not found |
| DGAP133 | 6q13-q14.1 deleted includes the epigenetic controller **HMGN3** which normally stimulates acetylation of histone H3. <sup>26</sup> | SMAP1, B3GAT2, OGFRL1, RIMS1, SLC17A5, FILIP1, LCA5, SENP6, etc. (See Fig 1) | KHDC1, EEFA1, MYO6, IRAK1BP1, B3GAT2, etc. (see Fig 1) | Collagen genes 19A, 12A, COX7A2 | Col9A |
| DGAP139 | 13q14.2 deleted Includes SETDB2 a gene that codes for the epigenetic regulator methyltransferase <sup>27</sup> (See Fig. 1) | SUCLA2, ITM2B, RCBTB2, CYSLTR2, KPNA3 | RBI, NUDT15, DLEU1,2, PHF11, SETDB2, KCN-RG | ITM2B, CYSLTR2, LPAR6, DLEU2 | DLeu2 |
| DGAP142 | **MBD5** Epigenetic regulator Histone acetylation disrupted by rearrangement. | ✓ | ✓ | ✓ | Not found |
| DGAP145 | **EFTUD2** [RNA splicing regulator in spliceosome] at rearrangement breakpoint. An epigenetic effector that connects methylated histone to RNA splicing <sup>28</sup> | ✓ | ✓ | ✓ | ✓ |
| DGAP147 | NALCN encodes sodium leak channel | ✓ | ✓ | Not found | Not found |
| DGAP154 | Xq25 (Duplication) XIAP, **STAG2** <sup>29</sup>. STAG2 inhibition increase the ability to reprogram cardiac fibroblasts implicating STAG2 in epigenetics. | XIAP, THOC2, GRIA3 | XIAP | SMAD5 | SMAD5 |
| DGAP155 | **EHMT1**/GLP: Encodes a histone methyltransferase, an epigenetic regulator. | ✓ | ✓ | ✓ | ✓ |
| DGAP157 | **FOXPI** "PRMT5 recruitment to the FOXPI promoter facilitates H3R2me2s, SET1 recruitment, H3K4me3, and gene expression" <sup>30</sup> Disrupted by rearrangement. | ✓ | ✓ | ✓ | ✓ |
| DGAP159 | 10p15.3-p14 Deletion includes **GATA3** See Fig. 1. **GATA3** controls enhancer accessibility,<sup>31</sup> | ✓ | ✓ | ✓ | ✓ |
| DGAP161 | 4q34 (See Fig. 1) **HMGB2**, effects epigenetic status<sup>32</sup> | AGA, SPCS3, TENM | HMGB2, HPGD, VEGFc ADAM29 | VEGFc, ADAM29 | HMGB2, VEGFc |
| DGAP164 | **NODAL/activin a** targets KDM6B (epigenetic regulator of neuronal plasticity) signaling in mouse brain. Rearrangement also disrupts **TET1** (a methylation eraser (demethylase)) | ✓ | ✓ | ✓ | ✓ |
| DGAP166 | **SCN1A** Sodium voltage gated channel | ✓ | ✓ | Not found | Not found |
| DGAP169 | **NR2F1** disrupted by rearrangement, target of the epigenetic regulator complex PRC2 | ✓ | Not found | ✓ | ✓ |
| DGAP173 | 11p14.2 FIBIN, BBOX1, SLC5A12, ANO3 | ANO3, BBOX1 | Mucin15 | Not found | Not found |
| DGAP186 | **NR5A1** disrupted, participates in resetting pluripotency. Differentially methylated in endometriosis | ✓ | ✓ | ✓ | Not found |
| DGAP189 | **SOX5** participates in epigenetics as a histone demethylase | ✓ | ✓ | ✓ | ✓ |
| DGAP190 | **SMS** Spermine synthetase gene. Polyamines control the epigenetic regulators histone demethylases | ✓ | ✓ | ✓ | ✓ |
| DGAP193 | SPAST | ✓ | ✓ | ✓ | ✓ |
| DGAP199 | NOTCH2, disrupted at breakpoint by rearrangement. | ✓ | ✓ | ✓ | ✓ |
| DGAP201 | **AUTS2** Associated with H3K4me3 and epigenetic control | ✓ | ✓ | Not found | Not found |
| DGAP202 | **KDM6A** Histone di/tri demethylase disrupted at breakpoint by rearrangement. | ✓ | ✓ | ✓ |  |
| DGAP211 | **SATB2** Epigenetic functions, interacts with chromatin remodelers. | ✓ | ✓ | Not found | ✓ |
| DGAP219 | **CUL3** Ubiquitin ligase recruited to epigenetically regulate lymphoid effector programs. Disrupted at breakpoint of rearrangement. | ✓ | ✓ | ✓ | ✓ |
| DGAP232 | **SNRPN-SNURF** / SMN; PWCR; SM-D; sm-N; RT-LI; HCERN3; SNRNP-N; Required for epigenetic imprinting. | ✓ | ✓ | Not found | ✓ |
| DGAP235 | **MBD5** Epigenetic regulator involved in histone acetylation disrupted at breakpoint of rearrangement. | ✓ | ✓ | ✓ | Not found |
| DGAP239 | **CHD7** Chromodomain helicase that collaborates with the epigenetic regulator SOX2<sup>33</sup> CHD7 is gene that encodes functions essential for chromatin remodeling | ✓ | ✓ | Not found | ✓ |
| DGAP244 | CTNND2 | ✓ | ✓ | Not found | ✓ |
| DGAP278 | **SNRPN-SNURF** / SMN; PWCR; SM-D; sm-N; RT-LI; HCERN3; SNRNP-N; Required for epigenetic imprinting | ✓ | ✓ | Not found | ✓ |
| DGAP301 | **MEF2C**, Associated with histone hypermethylation in disease<sup>34</sup>.. Disrupted by rearrangement | ✓ | ✓ | ✓ | ✓ |
| DGAP316 | 18p11.32-p11.22 deletion: **PHF21A** "Histone methyl-reader"<sup>20</sup> within the histone deacetylase complex PHF21A Encodes BHC80, a component of a BRAF35 histone deacetylase complex | ✓ | ✓ | YES1, MYOM1, EPB41L3, ARHGAP28, LAMA1, PTPRM, MTCL1, TWSG1, RALBP1, RAB31 | SMCHD1, TGIF1, LAMA1, PTPRM, TWSGH1, |
| DGAP317 | 6q14.1 TBX18: IRAK1BP1; IBTK | LCA5, ELOV4, HTR1B, DOPEY | IRAK1BP1, TBX18 IBTK NFKB | COX7A | Not found |
| MGH7 | **GRIN2B** Reflects changes in methylation levels<sup>35</sup> | ✓ | ✓ | Not found | Not found |
| MGH8 | **CHD8**, CHD8 interacts with the insulator binding protein CTCF to insulate and epigenetically regulate<sup>36</sup>. | ✓ | ✓ | Not found | Not found |
| MGH9 | **TCF4** regulated by histone deacetylases. Binding sites overlaps histone acetylation of enhancers<sup>37</sup> | ✓ | ✓ | ✓ | ✓ |
| NIJ2 | **PHIP**, linked to histone methylation <sup>38</sup> Disrupted at breakpoint by rearrangement. | PHIP, MYO6 | PHIP, MYO6 | PHIP, MYO6 | PHIP, MYO6 |
| NIJ5 | **ILIRAPL1** | ✓ | ✓ | Not found | Skeletal growth |
| NIJ6 | **KAT6B** / MORF/ MYST4 Lysine acetyl transferase. | ✓ | ✓ | ✓ | ✓ |
| NIJ14 | **NFIA**, regulated by chromatin structure <sup>39</sup> | ✓ | ✓ | ✓ | ✓ |
| NIJ15, NIJ6 | **MYT1L** | ✓ | ✓ | Not found | Not found |
| ROC4 | **CAMTA1** | ✓ | ✓ | ✓ | ✓ |
| ROC17 | **AUTS2** Associated with H3K4me3 and epigenetic control | ✓ | ✓ | Not found | Not found |
| ROC23 | **TCF12** | ✓ | ✓ | ✓ | ✓ |
| ROC43 | **SOX5** is a histone demethylase | ✓ | ✓ | ✓ | ✓ |
| ROC62 | **SNRPN-SNURF** Required for epigenetic imprinting | ✓ | ✓ | Not found | ✓ |
| UTR7 | **NFIX**, controlled by methylation | ✓ | ✓ | ✓ | ✓ |
| UTR12 | **DYRK1A**, histone phosphorylation | ✓ | ✓ | ✓ | ✓ |
| UTR13 | **MBD5**, Epigenetic regulator disrupted in topology associated domain | ✓ | ✓ | ✓ | Not found |
| UTR17 | **ZBTB20**, binds to genes that control chromatin architecture including MEF2c and SAT2b) | ✓ | ✓ | ✓ | ✓ |
| UTR20 | **FOXP2** | ✓ | ✓ | ✓ | ✓ |
| UTR21 | **NSD1** lysine methylation | ✓ | ✓ | ✓ | ✓ |
| UTR22 | 2q24.3 SCN9A | SCN9A, TTC21B, STK39 | NOSTRIN, CERS6 | FIGN, XIRP2, CERS6 | TTC21B |
| Affected member of family 1 | **NCALDI** exists in myristoyl and non myristoyl forms. Histone deacetylase 11 is a myristoyl hydrolase <sup>40</sup>. | ✓ | ✓ | Not found | ✓ |
| Affected member of family 1 | **NPL** - N-acetylneuraminatase lyases regulate cellular sialic acid concentrations by mediating its reversible conversion into N-acetylmannosamine and pyruvate | ✓ | ✓ | ✓ | ✓ |
| Affected member of family 1 | **BCAR3**. A candidate epigenetic mediator for increased adult body mass index in socially disadvantaged females <sup>41</sup> | Not found | Not found | ✓ | ✓ |
| Affected member of family 2 | **ZNF423** - epigenetic modifications associated with obesity <sup>42</sup>, methylation marker for age prediction. | ✓ | ✓ | Not found | ✓ |
| Affected member of family 3 | **RAP1GDS1** - **RAP1** GTP-GDP dissociation stimulator 1 <sup>43</sup>. Antagonizes some histones. | ✓ | ✓ | ✓ | ✓ |

**Figure 4.**
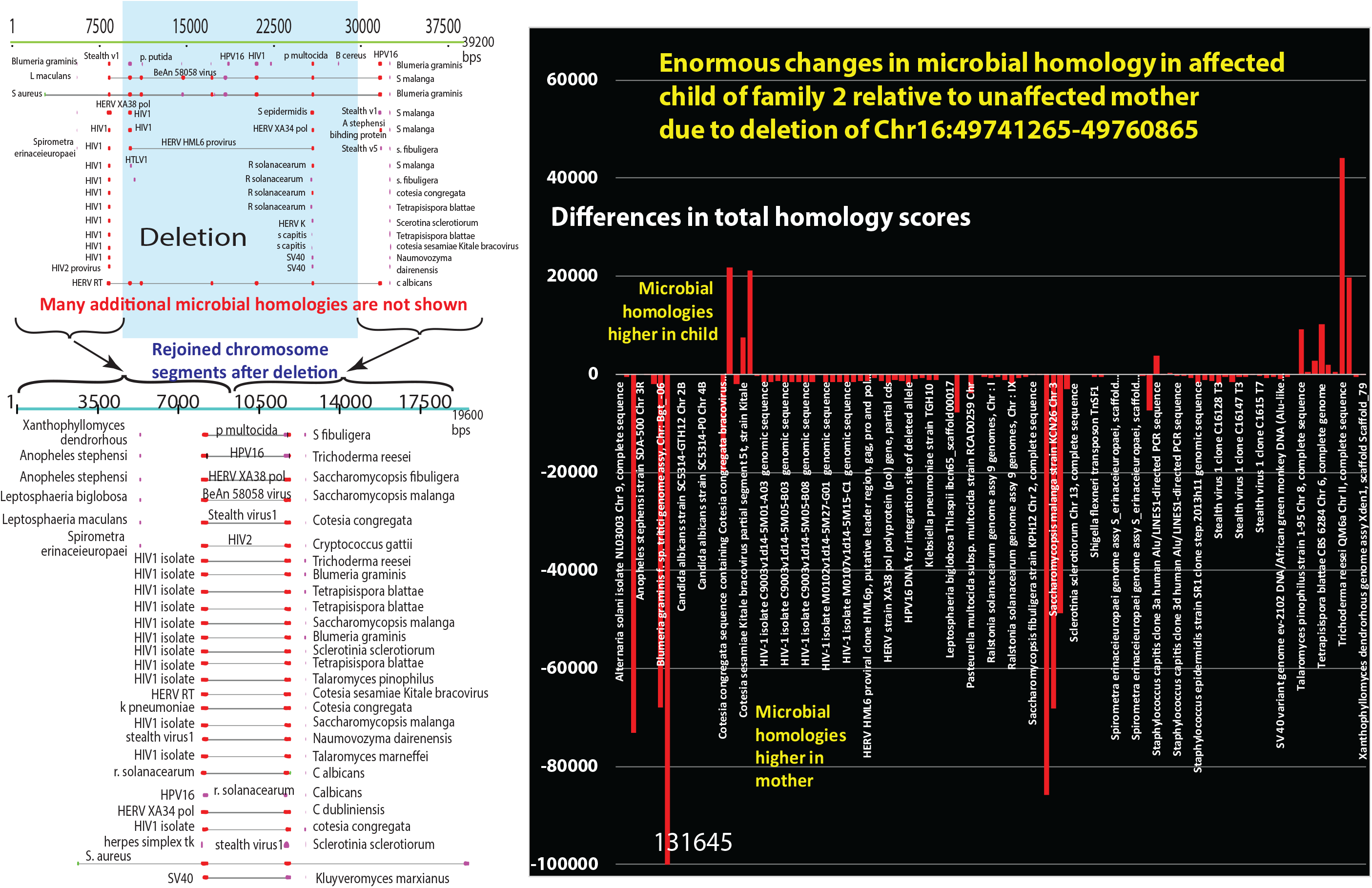
A structural variant unique to an affected member of family 2 in reference 3 has massive changes in the distribution and identities of homologous microorganisms when compared to the unaffected mother. At the left (top) are representations of the homologies to normal human chromosome 16:49741265-49760865 [GrCh37] with 9600 bps on both the 5’ and 3’ ends. After the deletion (lower left), the two 9600 bp additions are juxtaposed. The rearrangment creates multiple differences in the distributions of foreign DNA hoomologies. The graph at the right shows the quantitive differences created by the rearrangement. Microorganisms are arranged alphabetically. Bars above the line are microbes more strongly represented in the affected child’s sequence and those below the line are stronger in the unaffected mother’s sequence. This result was repeated many times with a variety of different assumptions both for the regions and for the homology criteria.

### Some chromosome regions with microbial homologies are only deleted in affected family members in families that share a recurrent translocation

Recurrent de novo translocations between chromosomes 11 and 22 have so far only been detected during spermatogenesis and have been attributed to palindromic structures that induce genomic instability^2^. The recurrent breakpoint t(8;22)(q24.13;q11.21) ^2^ was tested to determine whether palindromic rearrangements might arise because microbial infection interferes with normal chromatin structures.

Fig. 5 shows strong homology to bacterial and viral sequences in a family with a recurrent translocation. Sequences from an unaffected mother carry a balanced translocation rearrangement ^2^ with homologies to more diverse microbes than are present in affected cases (Fig. 5). The distributions of homologous microorganisms are clearly different for the unaffected mother vs affected Case 12 Der(8) (Mann-Whitney P=0.0049) and vs Case13 (not shown).The rearrangement accompanies profound local changes in the homologous microbiome (P=0.0022). Some prominent microbes that show large differences, e.g. fungi and yeasts have been isolated from oral cavities of children^14^.

**Fig. 5.**
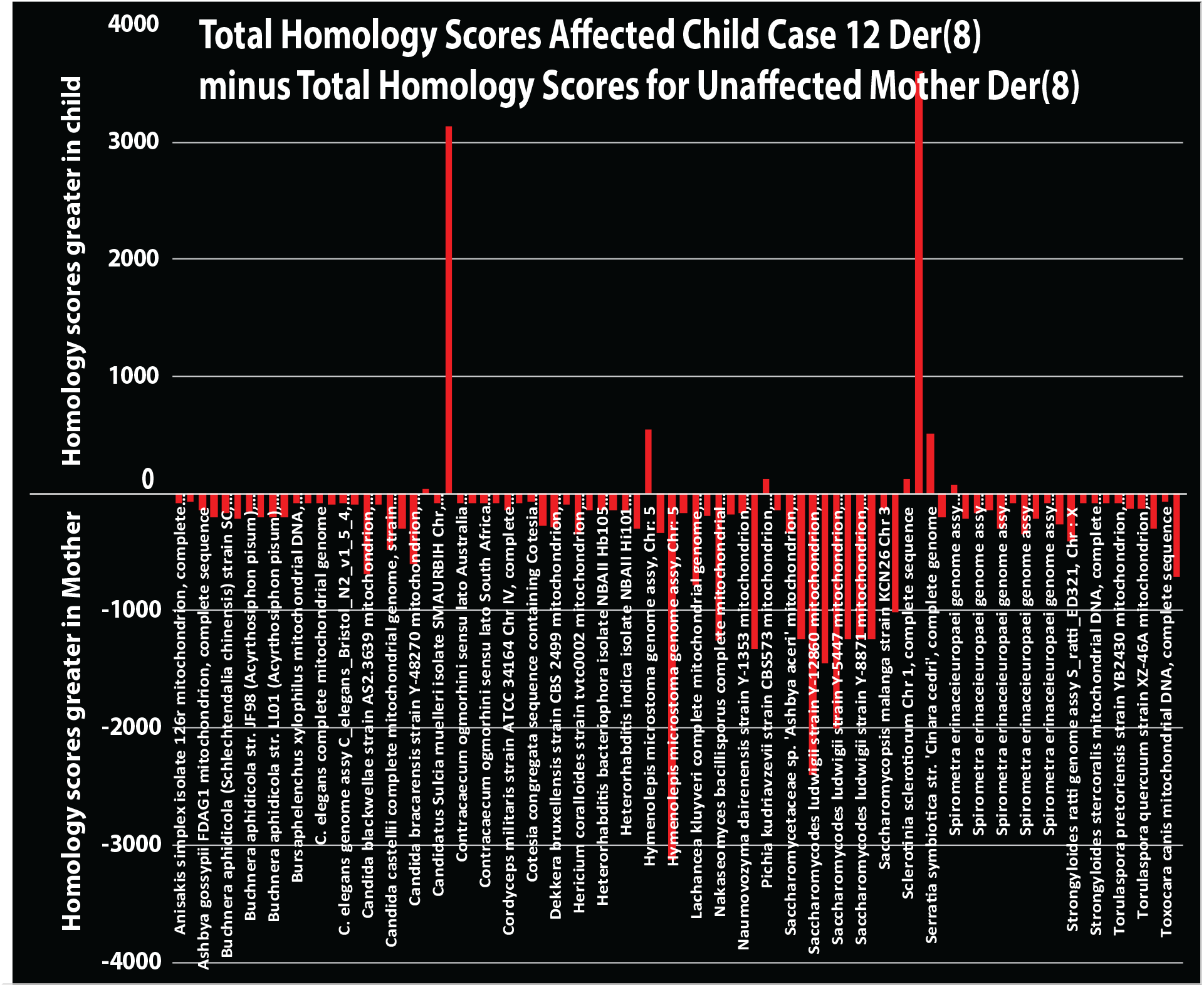
Changes in microbial homologies in affected child born to a mother with a recurrent translocation. Parental translocation t(8;22) in Family FHU13-027 from Mishra et al (reference 2) is homologous to microorganisms as shown for der(8). The graph represents the difference in total homology scores for an affected child Case 12 subtracted from those of the normal healthy mother with balanced translocation t(8;22). A very different set of micro-organisms is present in affected Case 12. Bars above the axis are mainly absent in the asymptomatic mother. Bars below the axis are present in the mother but limited or absent in her child. The homologous microbiome is in alphabetical order.

### Microbial DNA homologies in areas around a mutated epigenetic driver gene

In some patients with neurodevelopmental disorders, a chromosomal anomaly disrupts a critical driver gene and there is strong evidence that the disrupted driver contributes to the disease ^1^. The genes identified as underlying phenotypic drivers of congenital neurologic diseases include chromatin modifiers ^1^. In agreement with this designation,Table 1 shows that most pathogenic driver genes are more specifically epigenetic factors (at least 45 of the 66 patients listed in Table 1). Using a value of 815 as a rough estimate of the total number of epigenetic factors in the human genome^15^ containing 20,000 genes, the probability that association between neurodevelopmental patients and epigenetic modifications occurs by chance is P<0.0001.

### Pathogenic driver gene mutations caused by large chromosome deletions amplify their effects because of epigenetics

Because most identified driver genes of neurodevelopmental disorders ^1^ are epigenetic factors (Table 1), the functions they control in individual patients and in families with members affected by neurodevelopmental disorders ^3^ were compared to genes in pathogenic chromosomal deletions. Parts of pathogenic chromosome deletions affected these kinds of critical neurodevelopmental driver genes ^1^. Like clustered chromosomal deletions, virtually all pathogenic driver genes have strong effects on the immune system, angiogenesis, circulation and craniofacial development. Fig. 6 summarizes how the functions of damaged epigenetic drivers are distributed and shows that all 46 gene drivers of neurodevelopment have pleiotropic effects. By comparison, pleiotropy has been documented for 44% of 14459 genes in the GWAS catalog ^16^. By this standard, neurodevelopmental driver genes are disproportionally pleiotropic (P<0.0001).

**Fig. 6.**
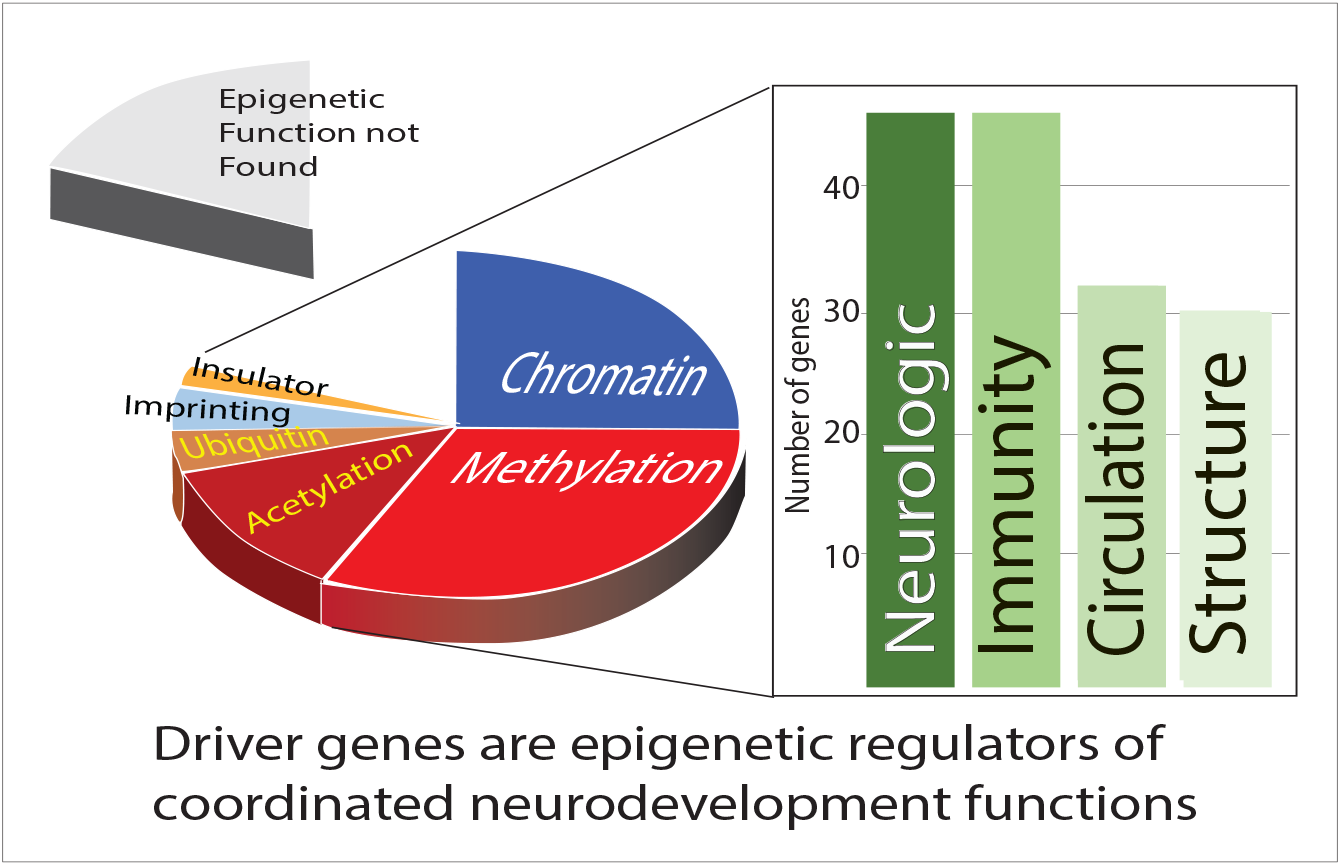
As a result of chromosome anomalies, driver genes truncated or deleted in congenital neurodevelopmental disorders are mainly epigenetic regulators or effectors. The pie chart shows the percentages of 46 driver genes that have the epigenetic functions indicated. Loss of these driver gene functions then impacts a group of functions that must be synchronized during the complex process of neurodevelopment. These are the same general functions lost in deleted gene clusters.

### Clear evidence of a non-human insertion

In 48 patients^1^, multiple infection matching sequences were included in chromosomal anomalies generated by balanced chromosomal translocations (Data not shown). Sequences around individual breakpoints were tested for microbial insertions by first comparing the sequences to human and then to microbial DNA. For example, chromosome breakpoint 2 in patient DGAP154 matched human DNA X-chromosome in two segments with a gap in the sequence (Fig. 7). The gap did not match human sequences but did correspond to nematodes and yeast-like fungi, suggesting one or more of these microorganisms had inserted foreign DNA into patient DGAP154 (Fig. 7). More frequently however, other breakpoints in DGAP154 chromosomes matched many microbial sequences. A simple example of one of these alignments around DGAP154 breakpoint 3 shows that many microorganisms align with human DNA. The similarity between critical human DNA epigenetic factors and microbial DNA (which may be more abundant) can set up competitions during recombination and break repair (Fig 7 bottom).

**Fig. 7.**
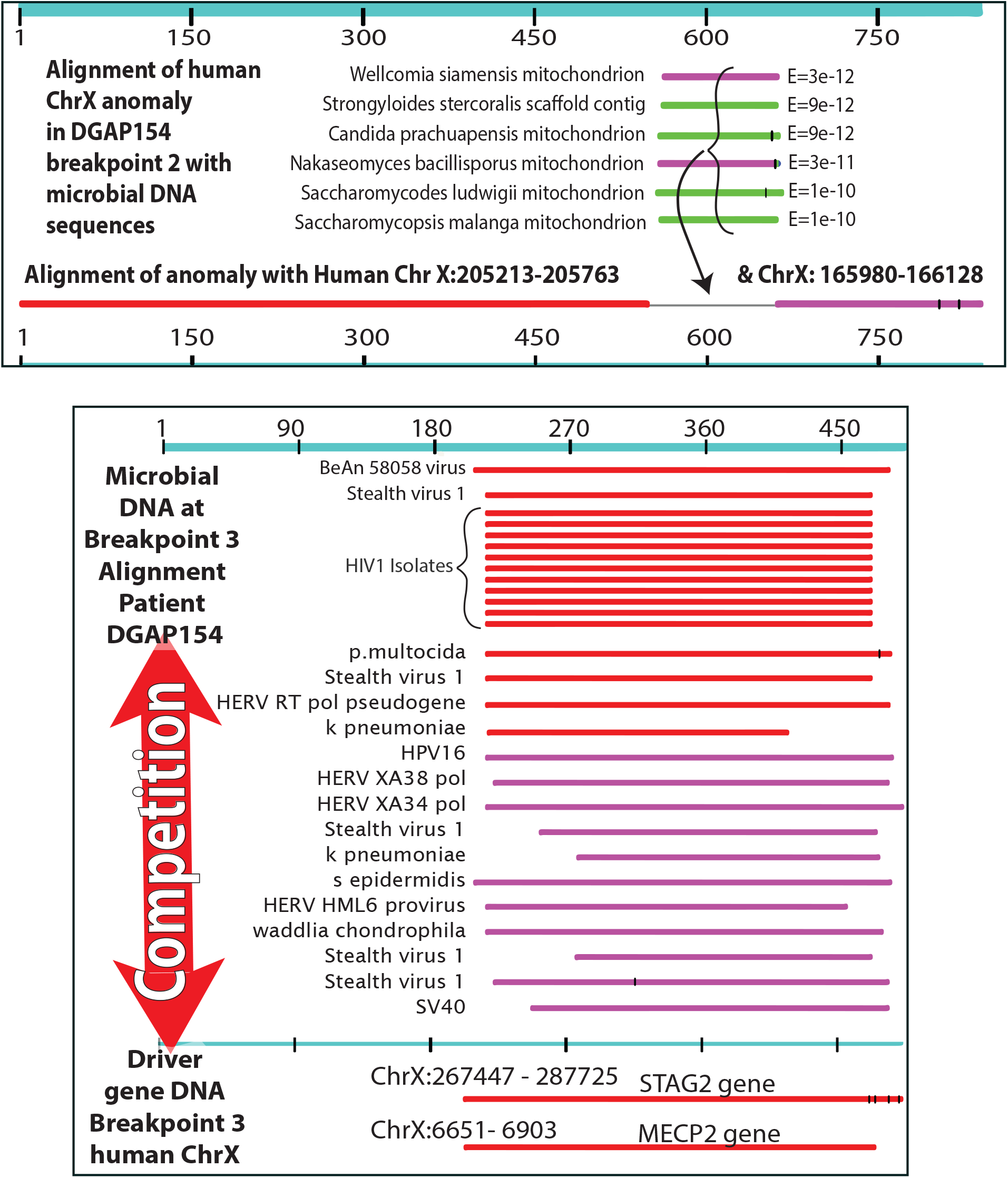
Foreign sequences can compete with human DNA at epigenetic regulators around breakpoints. In the top panel Breakpoint 2 in patient DGAP154), Mitochondrial DNA from potential pathogensnematodes and yeast-like fungi, aligns well with an anomaly in a damaged epigenetic factor. The sequence at Breakpoint 2 on chromosome X indicated by the black line does not match any human sequences. Any of the nemotodes or yeast-like fungi could insert their DNA at this position. The lower panel (Breakpoint 3 in the same patient) may be more typical and shows that microbial DNA can compete with human DNA around damaged epigenetic regulators such as MECP2. Microbial homology to this major epigenetic regulator was confirmed by testing human MECP2 against microbes. STAG 2 is also implicated in epigenetics (Table 1).17

### Multiple infections identified by homology match signs and symptoms of neurodevelopmental disorders

These kinds of alignments suggest candidates that can contribute to the signs and symptoms in each individual (Table 2). 8 patients have growth retardation. 20 of 48 patients had impaired speech ^1^. Multiple infections can cause these problems. For instance, HIV-1 causes white matter lesions associated with language impairments and impaired fetal growth. There are nearly 50 matches to HIV-1 DNA in the chromosome anomalies of 35 patients.

**Table 2.**
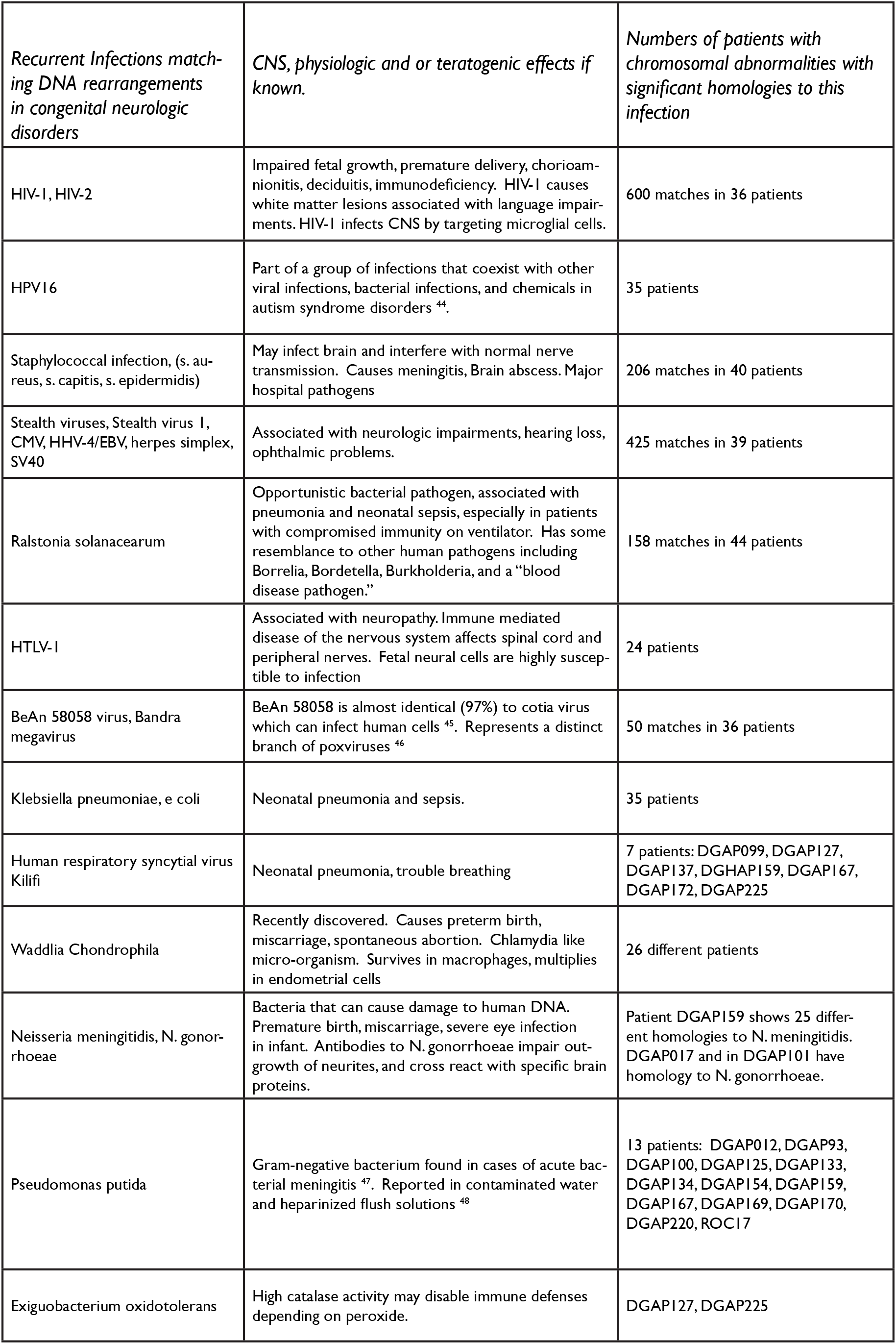
Recurrent infections found to have homology to chromosomal abnormalities in neurologic birth defects can cause developmental defects.

| *Recurrent Infections matching DNA rearrangements in congenital neurologic disorders* | *CNS, physiologic and or teratogenic effects if known.* | *Numbers of patients with chromosomal abnormalities with significant homologies to this infection* |
| --- | --- | --- |
| HIV-1, HIV-2 | Impaired fetal growth, premature delivery, chorioamnionitis, deciduitis, immunodeficiency. HIV-1 causes white matter lesions associated with language impairments. HIV-1 infects CNS by targeting microglial cells. | 600 matches in 36 patients |
| HPV16 | Part of a group of infections that coexist with other viral infections, bacterial infections, and chemicals in autism syndrome disorders <sup>44</sup>. | 35 patients |
| Staphylococcal infection, (s. aureus, s. capitis, s. epidermidis) | May infect brain and interfere with normal nerve transmission. Causes meningitis, Brain abscess. Major hospital pathogens | 206 matches in 40 patients |
| Stealth viruses, Stealth virus 1, CMV, HHV-4/EBV, herpes simplex, SV40 | Associated with neurologic impairments, hearing loss, ophthalmic problems. | 425 matches in 39 patients |
| Ralstonia solanacearum | Opportunistic bacterial pathogen, associated with pneumonia and neonatal sepsis, especially in patients with compromised immunity on ventilator. Has some resemblance to other human pathogens including Borrelia, Bordetella, Burkholderia, and a "blood disease pathogen." | 158 matches in 44 patients |
| HTLV-I | Associated with neuropathy. Immune mediated disease of the nervous system affects spinal cord and peripheral nerves. Fetal neural cells are highly susceptible to infection | 24 patients |
| BeAn 58058 virus, Bandra megavirus | BeAn 58058 is almost identical (97%) to cotia virus which can infect human cells <sup>45</sup>. Represents a distinct branch of poxviruses <sup>46</sup> | 50 matches in 36 patients |
| Klebsiella pneumoniae, e coli | Neonatal pneumonia and sepsis. | 35 patients |
| Human respiratory syncytial virus Kilifi | Neonatal pneumonia, trouble breathing | 7 patients: DGAP099, DGAP127, DGAP137, DGHAP159, DGAP167, DGAP172, DGAP225 |
| Waddlia Chondrophila | Recently discovered. Causes preterm birth, miscarriage, spontaneous abortion. Chlamydia like micro-organism. Survives in macrophages, multiplies in endometrial cells | 26 different patients |
| Neisseria meningitidis, N. gonorrhoeae | Bacteria that can cause damage to human DNA. Premature birth, miscarriage, severe eye infection in infant. Antibodies to N. gonorrhoeae impair outgrowth of neurites, and cross react with specific brain proteins. | Patient DGAP159 shows 25 different homologies to N. meningitidis. DGAP017 and in DGAP101 have homology to N. gonorrhoeae. |
| Pseudomonas putida | Gram-negative bacterium found in cases of acute bacterial meningitis <sup>47</sup>. Reported in contaminated water and heparinized flush solutions <sup>48</sup> | 13 patients: DGAP012, DGAP93, DGAP100, DGAP125, DGAP133, DGAP134, DGAP154, DGAP159, DGAP167, DGAP169, DGAP170, DGAP220, ROC17 |
| Exiguobacterium oxidotolerans | High catalase activity may disable immune defenses depending on peroxide. | DGAP127, DGAP225 |

Within chromosome anomalies, stealth viruses have about 35 matching sequences. Stealth viruses are mostly herpes virus derivatives that emerge in immunosuppressed patients such as cytomegalovirus (CMV). Stealth virus 1 (Table 2) is Simian CMV with up to 95% sequence identity to isolates from human patients. First trimester CMV infection can cause severe cerebral abnormalities followed by neurologic symptoms ^49^. CMV is also a common cause of congenital deafness and visual abnormalities. 27 of 48 neurodevelopmental patients had hearing loss. Herpes simplex virus is another stealth virus that directly infects the central nervous system and can cause seizures (reported for 9 patients).

Chromosome anomalies in patient DGAP159 have strong homology to N. meningitidis. Signs and symptoms in patient DGAP159 are consistent with known neurodevelopmental effects of bacterial meningitis including hearing loss, developmental delay, speech failure and visual problems.

### Tests for artifacts in matches to human-microbial DNA

Genome rearrangements for patient data ^1^ produced 1986 matches with E<=e-10 and a mean value of 83% identity to microbial sequences (range 66-100%).About 190 Alu sequences resembled microbial sequences, supporting the idea that homologies among repetitive human sequences and microbes are real. Correspondence between microbial sequences and multiple human repetitive sequences increases possibilities that microbial sequences can interfere with essential human processes. Contamination of microbial DNA sequences by human Alu elements ^50^, was ruled out by comparing about 450 AluJ,AluS and AluY sequences to all viruses and bacteria in the NCBI database. In contrast, LINE-1 elements (NM_001330612, NM_001353293.1, NM_001353279.1) had no significant homology to bacteria and viruses.

### Tests for DNA sequence artifacts

To further test the possibility that some versions of these microbial sequences were sequencing artifacts or contaminated by human genomes, microbial genomes were tested for homology to human genomes. Homologies to human sequences were found across multiple strains of the same microorganism (Table 3). For example, an Alu homologous region of the HIV-1 genome (bps 7300-9000) in 28 different HIV-1 isolates was compared. All 28 HIV sequences matched the same region of human DNA, at up to 98% identity. In contrast, only 1 of 20 zika virus sequences matched humans and was not considered further.

**Table 3.**
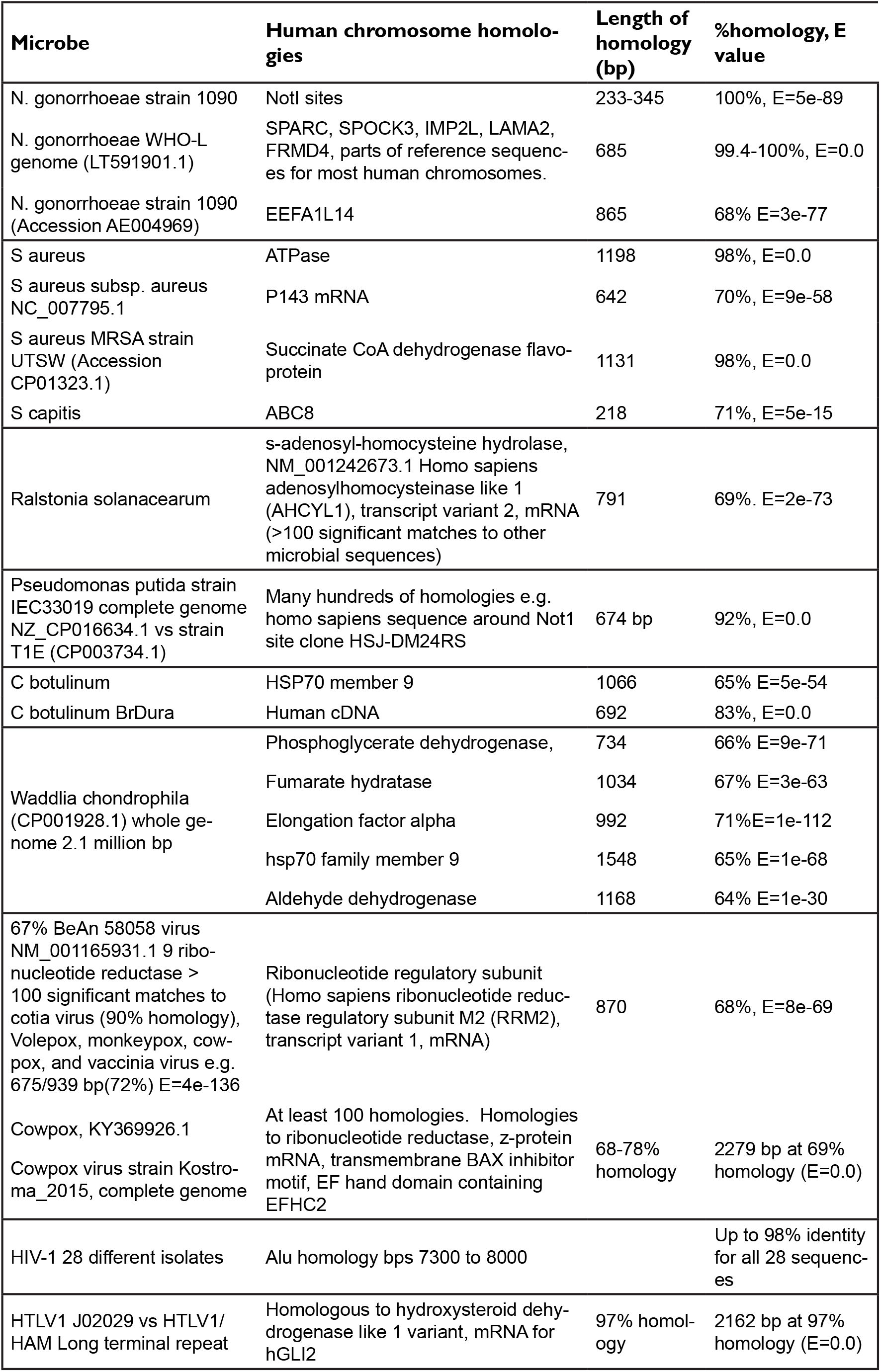
Independent evidence that microbial genomes have regions of homology to human DNA as predicted by results.

### A model for infection interference in neurodevelopment

Autosomal dominant inheritance of neurodevelopmental disorders containing microbial DNA suggest interference with gamete generation in one parent. The mechanism proposed in Fig. 8 is based on significant changes in microbial homologies on multiple human chromosomes. Large amounts of foreign DNA present during human meiosis with its many double strand breaks during the most active period of recombination produce defective gametes. Errors in spermatogenesis underlie a prevalent and recurrent gene rearrangement that causes intellectual disability, and dysmorphism (Emanuel syndrome) ^2^. In contrast, recombination in ova occurs in fetal life and then meiosis is arrested until puberty ^51^.

**Fig. 8.**
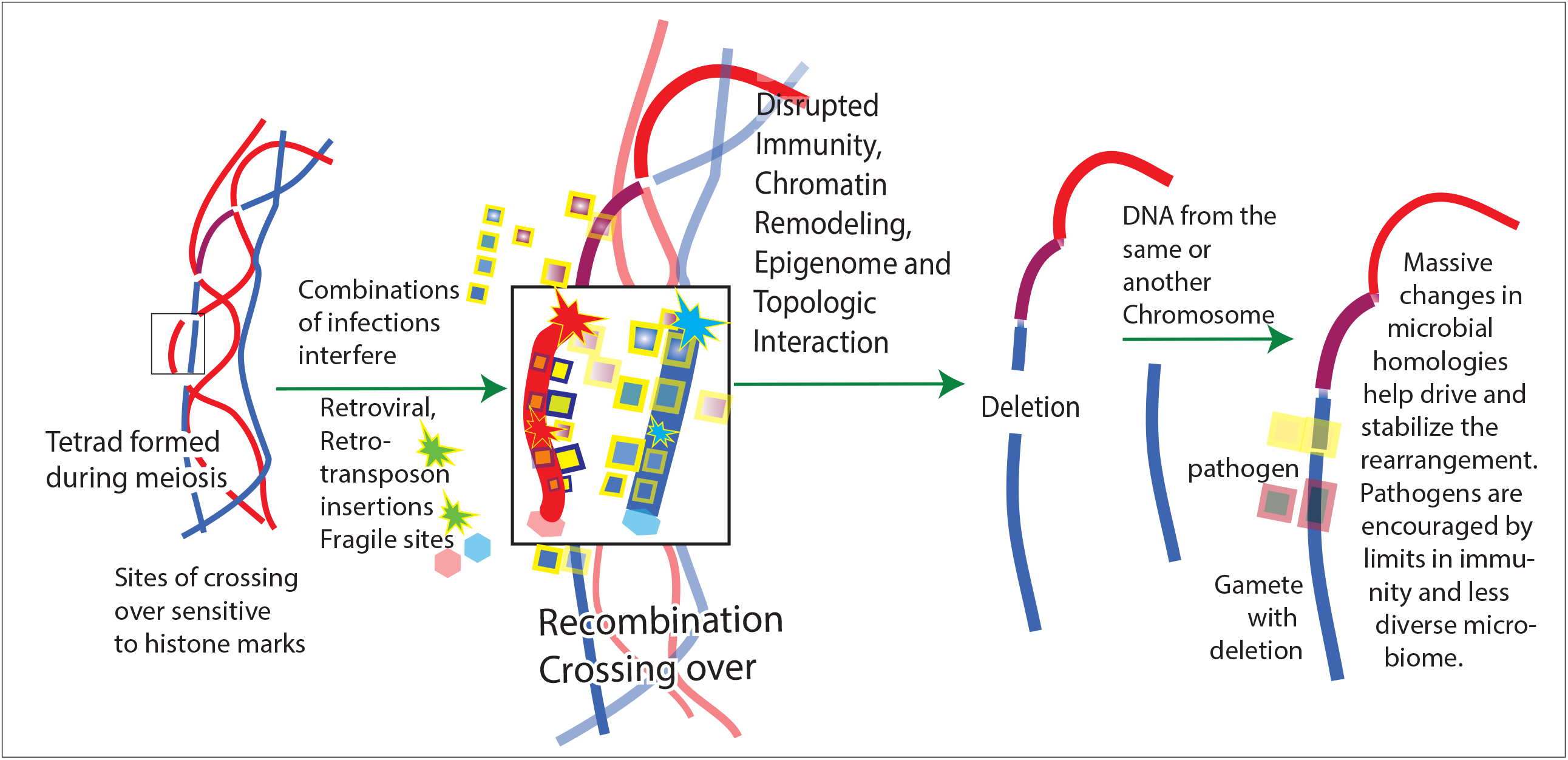
Soon after conception, erasure of epigenetic marks generates pluripotent stem cells and then the epigenome is reprogrammed. Foreign DNA can interfere with both processes leading to neurodevelopmental disorders. The model above shows interference with meiosis by DNA from infections. After duplication of parental chromosomes prior to generation of gametes, DNA from infections associates with strands of DNA in the many places including repetitive sequences of human DNA closely matching microbial DNA. In addition, retroviruses, retrotransposons may integrate their DNA and fragile sites may further destabilize the area. These events interfere with reductive cell division, topological relationships among chromosomes, epigenetic regulation and high-fidelity break repair. Interference from microorganisms may favor the illegitimate combinations due to palindromes reported by Mishra and co-workers ^2.^ Hundreds of DNA breaks occur during meiosis. Initiation sites for recombination are enriched in histone methylation and acetylation marks. Incorrect repair of recombination breaks is known to occur ^1^, causing chromosome anomalies such as deletions (shown). Massive changes in distributions of the homologous microbiome occurs to drive and stabilize the rearrangements. Clustered genes responsible for linked functions and epigenetic regulation in neurodevelopment are lost or displaced. Other chromosome segments with microbial homology do not contain identified genes but may be essential control regions, insulators, or essential for chromatin structures.

The resemblance of foreign microbial DNA to host background DNA may be a major factor in selecting infection and in human ability to clear the infection. Only one rare defective gamete is modeled in Fig. 8 but the male generates four gametes during meiosis beginning at puberty. Only one gamete survives in the female because three polar bodies are generated. Both balanced and unbalanced translocations can give rise to chromosome anomalies such as deletions and insertions because infection DNA can insert itself (e.g. exogenous or endogenous retroviruses), interfere with epigenomic marking or with break repairs during meiosis.A preexisting balanced chromosomal translocation in the family ^3^ increases the chances of generating a defective gamete during meiosis.

## Discussion

Long stretches of DNA in many infections match repetitive human DNA sequences which occur over a milion times. Individual microorganisms also match non-repetitive sequences. Human infections may be selected for and initially tolerated because of these matches. It is almost impossible to completely exclude the possibility of sequencing artifacts or contamination of microbial sequences with human sequences. However rather than reflecting widespread, wholesale error due to human DNA contamination in many laboratories over many years, microbial homologies more likely suggest that DNA sequences in the microbiome have been selected because they are homologous to regions of human DNA. This may be a driving force behind the much slower evolution of human repetitive DNAs.

Infections such as exogenous or endogenous retroviruses are known to insert into DNA hotspots^10^. Human DNA sequences that closely match infection DNA are proposed to generate chromosomal anomalies because similarities between microbial and human DNA interfere with epigenetic marking and with meiosis (Fig. 8). Thus, the genetic background of an individual may be a key factor in determining the susceptibility to infection and to the effects of infection. At the genetic level, this suggests selective pressures for infections to develop and use genes that are similar to human versions and to silence or mutate genes that are immunogenic. Infection genomes evolve rapidly on transfer to a new host ^52^. The presence of genes in infections that have long stretches of identity with human genes makes the infection more difficult to recognize as non-self. For example, there is no state of immunity to *N. gonorrhoeae*. Long stretches of *N. gonorrhoeae* DNA are almost identical to human DNA. Alu sequences and other repetitive elements are thought to underlie some diseases by interfering with correct homologous recombination as in hereditary colon cancer ^53^ or abnormal splicing. Why this does not always occur is not well understood. The presence of infection DNA that is homologous to multiple, long stretches of human DNA may mask proper recombination sites and encourage this abnormal behavior.

Neurons interact with cells in the immune system, sensing and adapting to their common environment. These interactions prevent multiple pathological changes ^54^. Many genes implicated in neurodevelopmental diseases reflect strong relationships between the immune system and the nervous system. It was always possible to find functions within the immune system for genes involved in neurodevelopmental disorders (Figs. 1 and 4). Damage to genes essential to prevent infection leads to more global developmental neurologic defects including intellectual disability. These homologies include known microbes knownt to produce teratogens. Analysis of mutations within clusters of genes deleted in neurodevelopmental disorders predict loss of brain-circulatory barriers, facilitating infections. Damage to cellular genes essential for autophagy may lead to abnormal pruning of neural connections during postnatal development.

Aggregated gene damage accounts for immune, circulatory, and structural deficits that accompany neurologic deficits. Other gene losses listed in Table 1 and in deleted chromosome segments (Fig. 1) account for deficits in cardiac function, cell barriers, bone structure, skull size, muscle tone and many other non-neurologic signs of neurodevelopmental disorders.

The arrangement of genes in clusters converging on the same biological process may simplify the regulation and coordination between neurons and other genes during neurodevelopment and neuroplasticity. Genes that are required for related functions, requiring coordinated regulation have been shown to be organized into individual topologically associated domains ^55^. Neurons are intimately connected to chromatin architecture and epigenetic controls ^56^. A disadvantage of the clustered arrangement of co-regulated accessory genes is that homology to microbial infections or other causes of chromosome anomalies anywhere in the cluster can then ruin complex coordinated neurological processes.

The results in Table 1 and Fig. 6 emphasize the role of epigenetic factors in neurodevelopmental diseases. Chromatin modifier genes are disproportionately affected in patients with neurodevelopmental disorders ^1^ and include two types of modifiers. Epigenetic factors signal chromatin remodelers, which are large multi-protein complexes. Epigenetic factors are responsible for differentiation from pluripotent states, chromatin remodeling also has major roles in developmental stage transitions.There are five families of chromatin remodelers that all control access to DNA within nucleosomes, exchanging and repositioning them. Chromatin remodeling arrays contain an ATPase subunit resembling motor proteins ^57^ and are distinct from epigenetic factors.

Epigenetic regulators that affect multiple functions required by the same process make their mutation especially critical. Longer range developmental interactions in chromosome regions exacerbate the effects of infection. Mutations or deletions (Fig. 1 and Table 1) show that this effect can occur in neurodevelopmental disorders. Functions that must be synchronized are grouped together on the same chromosome region and can be lost together.

Microbial DNA sequences are unlikely to be contaminants or sequencing artifacts. They are all found connected to human DNA in disease chromosomes; multiple microbial sequences from different laboratories are all homologous to the same Alu sequence. Alu element-containing RNA polymerase II transcripts (AluRNA’s) determine nucleolar structure and rRNA synthesis and may regulate nucleolar assembly as the cell cycle progresses and as the cell adapts to external signals ^58^. HIV-1 integration occurs with some preference near or within Alu repeats ^59^. Alu sequences are largely inactive retrotransposons, but some human-microbial homologies detected may be due to insertions from Alu or other repetitive sequences. Neuronal progenitors may support de novo retrotransposition in response to the environment or maternal factors ^60^.

Their variability and rarity make neurologic disorders difficult to study by conventional approaches. The techniques used here might eventually help predict the lifelong susceptibility to a given infection of an individual. However, a limitation is the inability to unequivocally identify one infection and to distinguish infection by one microorganism from multiple infections.

## Conclusions

1. DNAs in some congenital neurodevelopmental disorders closely matches multiple infections that extend over long linear stretches of human DNA and often involve repetitive human DNA sequences. The affected sequences are shown to exist as linear clusters of genes closely spaced in two dimensions.
2. Interference from infection can delete or damage human gene clusters and alter the epigenome. This interference accounts for immune, circulatory, and structural deficits that accompany neurologic deficits.
3. Neurodevelopmental disorders are proposed to begin when parental infections cause insertions or interfere with epigenetic markings and meiosis. Shifts in homologous microorganisms can be massive and may drive chromosomal rearrangements.
4. Congenital neurodevelopmental disorders are thus viewed as resulting from an assault on human DNA by microorganisms and an example of the selection of infecting microorganisms based on their similarity to host DNA.

## Acknowledgments

This work is based on the outstanding DNA sequence information and publications from investigators listed in the references. These results emphasize the importance of their work. It is a pleasure to acknowledge the inspiration, challenge and stimulation provided by the Department of Biochemistry and Molecular Genetics and the College of Medicine at UIC.

## Conflicts of Interest

The author declares no conflict of interest.

